# Identity and provenance of neighbors, genotype-specific traits and abiotic stress affect intraspecific interactions in the annual legume *Medicago truncatula*

**DOI:** 10.1101/2020.10.08.330944

**Authors:** Sara Tomiolo, Christian F. Damgaard, Sha Zhang, Simon Kelly, Ke Tao, Joëlle Ronfort, Lauréne Gay, Jean-Marie Prosperi, Simona Radutoiu, Bodil K. Ehlers

## Abstract

- Accounting for intraspecific variation may improve our understanding of species coexistence. However, our knowledge of what factors maintain intraspecific variation is limited. We predicted that 1) a plant grows larger when with non-kin (i.e. different genotypes) than kin (i.e. same genotype) neighbors, 2) abiotic stress alters the outcome of kin vs. non-kin interactions, 3) genetic identity of plants affects composition of soil microbiome.
- We set up mini-communities of *Medicago truncatula*, where focal genotypes were grown together with two kin or two non-kin neighbors from different origins. We analyzed how origin, identity of interacting genotypes and abiotic stress affected growth and fruit production. We also analyzed the composition of soil microbial communities.
- Focal plants grew larger in non-kin than in kin mini-communities. This pattern was stronger in low level of abiotic stress and when interacting genotypes were from similar origins. However, genotypic variation in growth and response to competition had a stronger effect on growth than mini-community type. Plant genotype identity did not affect soil microbiome.
- We find that intraspecific variation is affected by genotype-specific traits and abiotic stress. Geographic, rather than genetic, distance among interacting genotypes affects the outcome of intraspecific interactions.

## Introduction

Over the past decade, several studies uncovered that a large proportion of natural trait variation occurs within species (Siefert *et al*., 2015; Hart *et al*., 2017), and proposed that accounting for such diversity may improve our understanding of community-level processes (Vellend, 2010; Le Bagousse-Pinguet *et al*., 2014; Bolnick *et al*., 2011). Increasing numbers of theoretical and empirical studies have suggested that intraspecific genetic variation is likely to influence interactions between conspecifics, species richness, and ultimately species coexistence (Fridley & Grime, 2010; Jung *et al*., 2010; Ehlers *et al*., 2016). Although over the years many studies claimed the importance to understand the links between the behavior of individuals and the dynamics occurring at population- and community-level (Harper, 1977; Begon & Wall, 1987; Lichstein *et al*., 2007), we still lack an understanding of how genetic variation within species is maintained and how such variation may affect the outcome of intraspecific interactions.

We drew on species coexistence theory and studies at the interspecific and intraspecific level to formulate hypotheses about the factors that affect the growth and fitness of individuals growing with conspecifics. Two main hypotheses describe how plants may interact with conspecific neighbors based on their genetic relatedness. The niche partitioning hypothesis, which largely extrapolates from observations at the interspecific level, suggests that closely related individuals may have morphological similarities, similar phenotypes, and compete for the same niche (Young, 1981; Kelly, 1996; Burns & Strauss, 2012). For example, plants with similar heights will compete more strongly for light than plants of different height. Therefore, competitive exclusion is more likely to occur among closely related individuals, thus facilitating the coexistence of different and less closely related, genotypes. Kin selection theory, on the other hand, predicts that closely related conspecifics should divest from wasteful competition and redirect resources toward reproduction, thus increasing inclusive fitness (Hamilton, 1964a,b; West *et al*., 2002). To date, the debate between these two schools of thought has not yet been resolved (but see File *et al*., 2011).

Can species coexistence theory help predict what mechanisms maintain genetic variation within species? Species coexistence theory predicts that coexistence of different species can be mediated by different mechanisms (Aarssen, 1989; Chesson, 2000). Equalizing mechanisms, have been equated to a match between two boxers, in which neither is competitively superior to the opponent and where there is not a clear winner. In this case, coexistence of two species is maintained when none is competitively superior to the other (Aarssen, 1989). This is not a stable coexistence mechanism, as a shift in biotic or abiotic conditions may tip the scale and promote the success of one species over the other, thus leading to competitive exclusion. Stabilizing mechanisms ensure stable coexistence, by means of negative frequency-dependence, which favor rare species and limit the growth of abundant ones. In this case, no species can become too abundant, due to resource limitation, and no species can become too rare, because the decline of dominant species allows an increase of rare species (HilleRisLambers *et al*., 2012).

In addition, within a species some genotypes may be particularly favored due to a specific set of traits or due to favorable abiotic conditions, and become dominant (Fridley & Grime, 2010). In this scenario, if there is a sufficient difference in resource requirements among different genotypes, the increasing frequency of a genotype may be curbed once its pool of resources is depleted, thus favoring the increase of more rare genotypes, and ultimately allowing coexistence via stabilization (Le Bagousse-Pinguet *et al*., 2014). Some plant species that have strong small-scale genetic structure, occur in dense aggregates of siblings (i.e. kin) interspersed in a more heterogeneous matrix of distantly related conspecifics (i.e. non-kin). In this case, if closely related genotypes facilitate each other rather than competing, some genotypes may become increasingly abundant due to positive frequency dependence, until resources allow it (Ehlers & Bilde, 2019). If coexistence of kin is limited by resource availability, we would expect higher genotypic variation within species resulting from either equalizing mechanisms or niche differentiation, alternatively in the case of overlapping niches among different genotypes we would expect the dominance of the few stronger genotypes that have outcompeted the weaker ones.

Besides niche similarity and competition for shared resources, co-occurrence of different genotypes may be favored or hindered by abiotic and biotic factors (Gilman *et al*., 2010). The degree of abiotic stress experienced by individual plants may shift the outcome of their interactions with neighbors from positive to negative along a gradient of decreasing abiotic stress as predicted by the Stress Gradient Hypothesis (Bertness & Callaway, 1994). Finally, plants may increase their own fitness and the fitness of neighboring conspecifics by attracting favorable soil organisms in exchange for nutrients, and may thus ultimately influence the composition of soil biota (Reynolds *et al*., 2003; De Kroon *et al*., 2012; Van der Putten *et al*., 2013; Eisenhauer *et al*., 2017) and the outcome of plant-plant interactions among and within species (Laliberté *et al*., 2015; Siefert *et al*., 2018, 2019; Bever *et al*., 1997). While these mechanisms have received substantial attention at the interspecific level, evidence of their influence at the intraspecific level is still limited (but see Biswas & Wagner, 2014; Eränen & Kozlov, 2008; Fajardo & McIntire, 2011; Keller & Lau, 2018).

We used genotypes of the annual legume *Medicago truncatula* from different origins within its distribution range. Due to its high degree of selfing and its strong spatial genetic structure (Bataillon & Ronfort, 2006), this species occurs naturally both in clusters of genetically identical siblings produced by the same maternal individual, as well as in more heterogeneous matrix of closely and less closely related conspecifics (Bonnin *et al*., 2001; Siol *et al*., 2007). We set up small experimental mini-communities composed of three individuals belonging to either the same (*kin*) or to different (*non-kin*) genotypes. Because drought stress is an important component in the native distribution range of the focal species, we exposed the mini-communities to two water treatments. We tested how growth of focal plants and whole communities changed as a function of community type, water treatment, and genotype origins. In addition, we tested how the same explanatory variables affected fruit production of focal plants. Finally, we tested whether different genotype combinations and community types supported different soil microbial communities.

We hypothesized that: 1) If genetic relatedness is a proxy for niche similarity, plant growth may be reduced by limited resources when growing with kin, and may be influenced by genotype-specific traits when growing with non-kin genotypes, 2) Water availability may mediate the outcome of plant-plant interactions across mini-community types, 3) Non-kin mini-communities may be associated with a richer and more diverse soil microbial community than kin mini-communities.

## Materials and Methods

### Study species

*Medicago truncatula* is a widespread annual legume native to the Mediterranean basin. To represent the natural variation within the species in its distribution range, we used nine genotypes originating from natural populations located in the South of France, Algeria, Cyprus and Morocco (Supporting information - Table S1, Fig. S1). These genotypes belong to a core collection (CC16) maintained at INRA Montpellier (INRA *M. truncatula* Stock Center: www1.montpellier.inra.fr/BRC-MTR/), and were chosen because they capture the range of simple-sequence repeat (SSR) diversity found in a worldwide collection of naturally occurring lines (Ronfort *et al*., 2006). The seed lots used for this study were obtained after at least three successive generations of self-fertilization, and can thus be considered as homogeneous inbred lines.

*Medicago truncatula* reproduces predominantly through selfing (Bataillon & Ronfort, 2006). It produces spiny fruits (pods) containing multiple seeds, which fall to the ground when mature. When fruits fall close to the maternal plant, seeds will germinate and grow in dense clusters of kin. Alternatively, fruits can be passively dispersed by attaching to the hair of grazers or small mammals, and germinate into a more heterogeneous matrix of kin and non-kin genotypes. As a consequence, populations of *M. truncatula* are characterized by a strong genetic structure, where dense kin patches alternate with patches of different genotypes (Bonnin *et al*., 2001; Siol *et al*., 2008; Jullien *et al*., 2019).

### Greenhouse experiment

We set up mini-communities of three plants in a pot, where focal genotypes were exposed to either two neighbors of the same genotypes (which we refer to as *kin*) or of a different genotype (which we refer to as *non-kin*) originated from a different population. Seedlings were planted equidistant, so that each plant would have access to the same amount of resources and space within a pot. Because our genotypes originated from climatically different regions (Burgarella *et al*., 2016), non-kin treatments were mostly limited to genotypes of the same origin, except for the genotypes from Cyprus and Morocco, where climatic data (Burgarella *et al*. (2016); Supporting information - Table S2) suggested that genotypes had similar adaptations to drought. In total we built 25 different communities (Table 1). To check if the spatial distance between origins affected the degree of genetic relatedness among genotypes, calculated by the Rousset index (Rousset, 1997), we applied linear regression. We found no relationship between spatial distance among genotype combinations and their degree of genetic relatedness (F_1,70_ = 0.014, p = 0.903, R^2^ = −0.01, Supporting information - Fig. S2).

**Table 1:** accessions of *Medicago truncatula* used for this experiment

| Provenance | Accession | Combined with... |
| --- | --- | --- |
| Algeria | A_05 | A_5, A_11, A_14 |
| Algeria | A_08 | A_08, A_5, A_14 |
| Algeria | A_11 | A_11, A_5, A_14 |
| Algeria | A_14 | A_14, A_5, A_8 |
| France | F_07 | F_7, F_13, F_15 |
| France | F_13 | F_13, F_07, F_15 |
| France | F_15 | F_15, F_07, F_13 |
| Cyprus | C_02 | C_02, M_12 |
| Morocco | M_12 | M_12, C_02 |

Plants were initially germinated in trays. Subsequently, even-sized seedlings were transplanted in 5l pots (diameter = 23 cm, height = 17.9 cm) filled with a mixture of greenhouse soil: sand: vermiculite (1:1:1). Within each pot, the focal genotype was marked with a white plastic tag, while neighbor genotypes had a red or a green tag. In kin communities, where all genotypes were identical, tags were assigned randomly to each individual at the time of sowing. To test whether water availability affects the outcome of intraspecific plant-plant interactions, we exposed mini-communities to two water treatments: high, where plants were watered eight minutes per day, and low, where plants were watered two minutes per day by means of top irrigation. Throughout the experiment, we assessed the effectiveness of the two water treatments by regularly checking that soil in low water treatments was drier than in high water treatments, and that plants in the low water treatment showed signs of drought stress. In the low water treatments, we consistently observed that plant leaves were drooping and folded inward along their longitudinal axis (a sign of drought stress), whereas in the wet watering treatments plant leaves were fully open. The combination of community types (kin *vs*. non-kin) and water treatments (high *vs*. low) was replicated three times resulting in 150 pots in total (25 communities x two water treatments x three replicates). The position of pots within each water treatment was randomized regularly.

Starting one week after setting up the experiment, we monitored the growth of the focal and neighbor plants by measuring plant mean diameter and height for five weeks (four measurements in total). We also counted the number of leaves on each single plant, and we stopped after three measurements when, due to plant size, it became challenging to accurately count the total number of leaves on each individual plant.

When plants started to develop fruits, we bagged them singularly to avoid fruit loss and to correctly estimate reproductive output. Fruits were collected and counted at the end of the growing season in mid-October. Fruit number was used as an estimate of plant fitness.

### Soil microbiota sampling

When plants were 10 weeks old, we sampled the soil microbiota of each pot in the low water treatment - including three extra pots with no plants that were used as controls - for soil 16*S rRNA* v5-v7 amplicon sequencing. At this time, microbial communities would have had time to develop in pots. We used a plastic cylinder (2 cm diameter) to sample soil cores from the center of the pot, where we expected lower root concentration. We then selected a sample of soil from the middle of the soil core, manually removed any roots in the sample, and stored the soil samples in Eppendorf (1.5 ml) tubes. Tubes were immediately placed on dry ice and subsequently transferred to a freezer at −80°C degrees for further processing.

### Generation of 16S rRNA amplicon libraries for Illumina Miseq sequencing

At the end of the greenhouse experiment, we selected a subset of soil samples for 10 mini-communities (with three biological replicates for each) corresponding to pairs of kin and non-kin mini-communities. Our selection criteria were based on the strongest differences in focal genotype growth response between kin and non-kin mini-communities. The soil samples were homogenized using a Precellys Evolution (Bertin technologies). DNA was extracted using the FastDNA Spin kit for Soil (MP Bioproducts) according to the manufacturer’s protocol. DNA concentrations were measured fluorometrically (Quant-iT™425 PicoGreen dsDNA assay kit, Life Technologies, Darmstadt, Germany) and adjusted to 3.5 ng/*μ*l. The variable v5-v7 regions of 16S rRNA gene were amplified using primer pairs 799F and 1193R) (Chelius & Triplett, 2001; Bodenhausen *et al*., 2013). A second PCR was performed to incorporate barcode indexes and sequencing adapters. PCR products were purified, pooled in equal amounts and sequenced on an Illumina Miseq instrument by IMGM, Germany.

### Processing of 16S rRNA reads

All 16s rRNA amplicon reads were processed using QIIME (Caporaso *et al*., 2010) and USEARCH (Edgar, 2010) tool. Main steps for processing included merging of paired-end reads, quality filtering, de-replication, singletons and chimera removal, OTU clustering at a 97% threshold, generating OTU table, taxonomy assignment, and OTU table filtering. OTU clustering was conducted using the UNOISE method (Edgar, 2016). Taxonomy assignment was done with uclust (assign_taxonomy from QIIME) against the Greengenes (Greengenes OTUs 13_5) database (McDonald *et al*., 2012). OTUs that were assigned as mitochondrial or chloroplast and had relative abundance less than 0.01% were removed. The filtered OTUs table was used for downstream analysis.

### Statistical analyses

#### Generalized Birch model

We were interested in estimating the growth of whole mini-communities in response to water treatment and community type, as well as the growth of single genotypes while accounting for the growth of their neighbors. Thus, we chose a generalized Birch growth model (Damgaard & Weiner, 2008) because it allows 1) to estimate growth parameters both at the population (i.e. pot) and at the genotype level, 2) to estimate whether the growth of individuals is determined by non-competitive factors (such as genotype’s initial growth) or by the effects of neighbors’ growth later on in life (Damgaard & Weiner, 2008).

We used an estimate of plant volume in leaves equivalent (Supporting information - Fig. S3), which represented a good proxy for growth, to parameterize the degree of size-asymmetric growth in mini-communities (Supporting information Notes S1, Table S3). Two parameters describe the growth at the population (pot) level: *w* describes the global growth of plants within a pot, *a* describes asymmetry in growth within a pot (namely, how growth differs among individuals in the same pot). The first parameter, global growth at the pot level, was estimated as a response to water treatment as shown in the equation below (Eqn. 1). The second parameter, growth-asymmetry within a pot, was estimated in response to treatment, community type and growth of other individuals within the same pot.

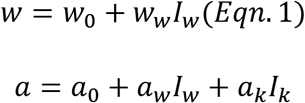

Where *I* is an indicator variable for water (*I_w_*) or community type (*I_k_*). *I_w_* equals 1 if the pot was exposed to high water treatment and 0 if the pot was exposed to low water treatment. *I_k_* equals 1 in kin mini-communities and 0 in non-kin mini-communities.

Two more parameters were estimated at the genotype level (Eqn. 2). The initial growth rate *r_i_*, which describes the growth rate of genotype *i* prior to experiencing competition from neighbors, and the inflection point of the growth curve *c_i_*, which describes the point at which the growth curve of genotype *i* slows and reaches the asymptote. The inflection point is determined by the effect of the growth of neighbor plants on the focal plant of genotype *i*. When a plant experiences strong competition from neighbors, the asymptote is reached earlier - and the value of *c_i_* is larger - compared to when a plant experiences weak competition from neighbors. The parameters *σ_r,i_* and *σ_c,i_* represent the variance of *r_i_* and *c_i_* respectively, the parameter *μ_r_* represents the mean growth rate and *μ_c_* the mean effect of neighbors on the growth curve of genotype *i*.

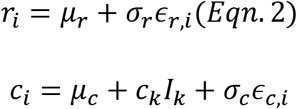

Based on the different combinations of the values of *r* and *c*, we predicted four theoretical growth trajectories of plant growth (Fig. 1A). In group 1 (green line), genotypes have a fast initial growth rate (*r*) and experience a low degree of competition (*c*) from neighbors; in group 2 (yellow line) genotypes have a fast initial growth rate, and experience strong competition from neighbors; in group 3 (red line), genotypes have slow initial growth rate, and experience low competition from neighbors; genotypes in group 4 (blue line) have a slow initial growth rate and experience strong competition from neighbors.

**Figure 1:**
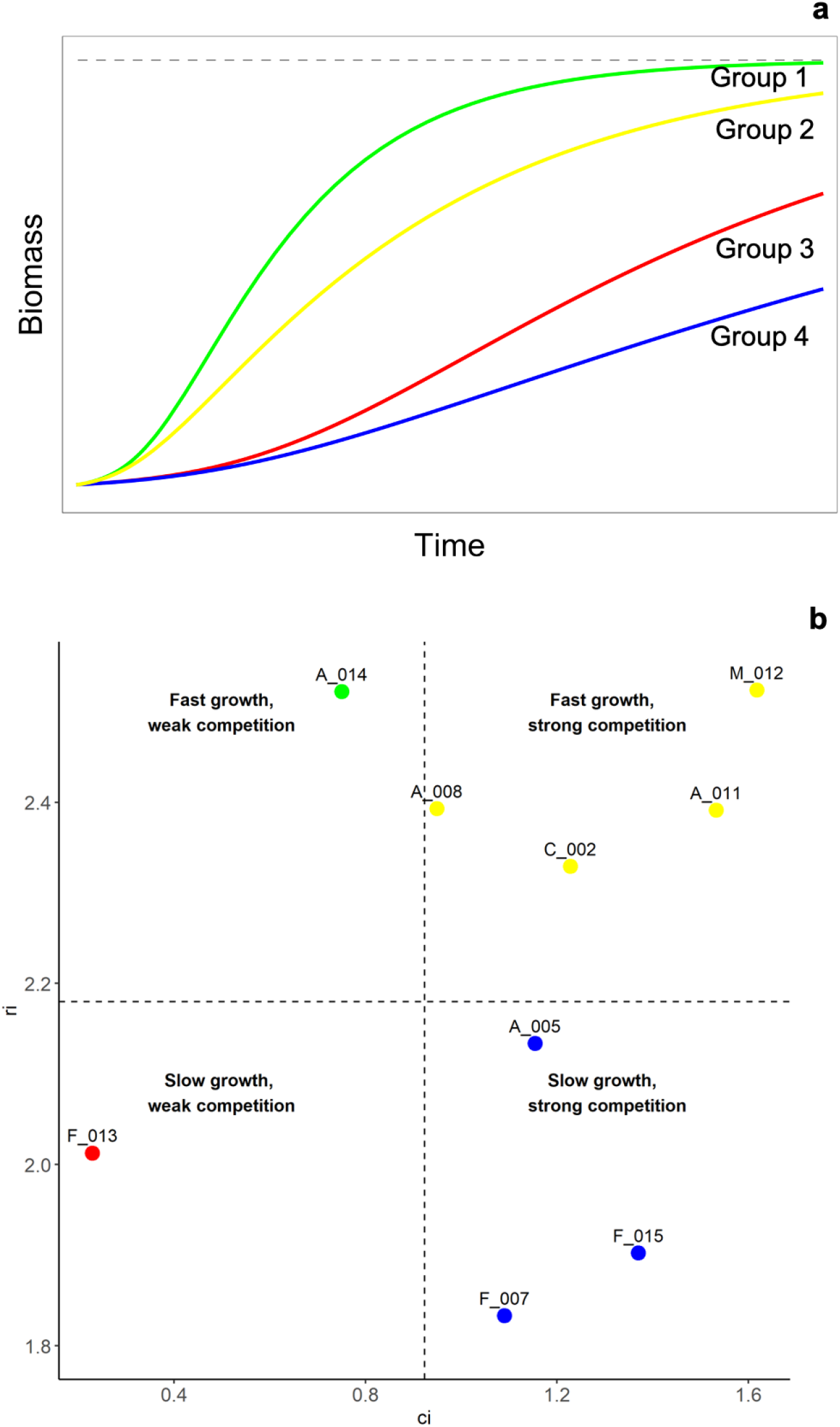
a) Conceptual representation of the Birch model, b)plot representing the ranking of genotypes based on the parameter estimates of the Birch model. a) The blue (group 4) and the red (group 3) curves represent genotypes experiencing initial slow growth rate (r_i_) and respectively strong and low competition from neighbors, the yellow (group 2) and green (group 1) line represent genotypes experiencing fast initial growth rate (r_i_) and respectively strong and low competition from neighbors. b) plot representing the ranking of genotypes based on the parameters r_i_ and c_i_ of the Birch growth model. Higher values of r_i_ indicate faster initial growth rate of a genotype, higher values of c_i_, indicate that the growth of neighbors has a stronger effect on growth of the focal. The cut-off lines are determined by the median values of c_i_ and r_i_, and the four resulting panels may be likened to the four strategies described in Fig. 1.

The joint Bayesian posterior distributions of the parameters were simulated by an MCMC approach using the Metropolis Hastings algorithm, assuming uniform improper prior distributions in the domains of the parameters. The MCMC iterations converged relatively fast after a burn-in period of 10000 iterations, and the next 40000 iterations were used to calculate the marginal posterior distribution and the corresponding 95% credibility interval of each parameter (Damgaard & Weiner, 2008). The model was developed using the software Mathematica (Wolfram, 2003).

#### Early life and fitness of focal plants

We did not carry out statistics on each single genotype combination due to the low number of replicates. We applied linear models to test the effects of water treatment, community type, region of origin, and their interaction on growth (measured as the increment in number of leaves) and on fruit production of the focal plants. Growth was measured as increment in number of leaves and was analyzed using linear models. Fruit production, calculated as number of fruits of focal plants, was analyzed using generalized linear models with negative binomial distribution using the R package MASS (Ripley *et al*., 2013). We selected the optimal models using the function *dredge* from the R package MuMIn (Barton & Barton, 2019). Significant interaction terms were tested using Tukey HSD posthoc tests in the R package emmeans (Lenth *et al*., 2019). These analyses were carried out in the software R3.6.1 (R Core Team, 2019).

#### Statistical analyses of soil microbiota composition

We characterized bacterial communities of unplanted soil and soil of kin and non-kin mini-communities, based on amplicon sequences from the V5-V7 hypervariable region of bacterial 16s rRNA gene. We rarefied the OTU table to the minimum read counts (single_rarefaction.py from QIIME) to calculate Shannon diversity indices among different mini-communities. Significant differences were determined using ANOVA *aov* function in R and post-hoc Tukey HSD test in R (p<0.05).

For calculating UniFrac distance (Lozupone & Knight, 2005) between samples, the OTU table was normalized using cumulative sum scaling (Paulson *et al*., 2013) and a phylogenetic tree was produced by the script (make_phylogeny.py from QIIME) (Price *et al*., 2010) according to the result of multiple sequence alignment generated by clustalo tool (Sievers *et al*., 2011; Sievers & Higgins, 2018). Weighted uniFrac distance was used as input for principle coordinate analysis (PCoA), using the *cmdscale* function in R (Anderson & Willis, 2003). Bray distances metric was used to calculate constrained analysis of principal coordinate (CPCoA), using the *capscale* function, in vegan package in R (Oksanen & others, 2015).

Finally, analyses of taxonomic composition profile were carried out to compare soil microbial community composition, among all mini-communities and among control, kin and non-kin communities. For this analysis, we focused on the relative abundance of the 15 top bacterial orders. The significance of difference in relative abundance was tested by Krustal-Wallis test (p<0.05).

## Results

### Birch’s growth model

Parameter estimates (Table 2,3) are described by the median of the posterior distributions and credibility intervals (2.5 and 97.5 percentiles). The estimated global plant growth at the pot level, which was calculated as *w* = *w*_0_ + *w_w_I_w_*, was significantly higher in the high water treatment (234.246 cm^3^) compared to the low water treatment (82.537 cm^3^). Within-pot size asymmetry, which was calculated as, *a* = *a*_0_ + *a_w_I_w_* + *a_k_I_k_*, was highest in non-kin mini-communities exposed to low water treatment (0.642) and lowest in kin communities exposed to high water (0.566).

**Table 2:** Estimate of the posterior distribution of the models parameters with prior according to Birch model, where 50% represents the median and 2.5% and 97.5% the credibility interval for the parameters. The parameters a = asymmetric growth and w = total pot biomass were estimated at the pot level, the parameters *μr, σr, μc, μck,σc*, were estimated at the genotype level.

| Parameter | 2.5% | 50% | 97.5% | P(X > 0) |
| --- | --- | --- | --- | --- |
| a0 | 0.56781279 | 0.642098655 | 0.716150069 | 1 |
| aw | -0.07121547 | -0.025505654 | 0.016096289 | 0.134569505 |
| ak | -0.089490204 | -0.050872252 | -0.015111855 | 0.003519546 |
| w0 | 60.99878172 | 82.53701508 | 131.9801737 | 1 |
| ww | 73.72192372 | 151.708823 | 216.5588468 | 1 |
| $\mu r$ | 2.030890389 | 2.288207483 | 2.511563839 | 1 |
| $\sigma r$ | 0.161060463 | 0.290363216 | 0.489506995 | 1 |
| $\mu c$ | 0.493172353 | 1.063149785 | 1.913964934 | 1 |
| $c_k$ | -0.558957681 | -0.061134465 | 0.222801865 | 0.303200661 |
| $\sigma c$ | 0.240796835 | 0.534515746 | 1.033260888 | 1 |
| $\sigma$ | 0.244185818 | 0.259941436 | 0.276104288 | |
| $\nu$ | 3.987539187 | 5.080487884 | 6.542017301 | |

**Table 3:** Values of *r* (initial growth rate of genotypes) and *c* (effect of growth from neighboring genotypes within a pot) calculated for each genotype *i*, where *r_i_* was calculated as *r_i_* = *μ_r_* + *σ_r_ϵ_r,i_* and *c_i_* was calculated as *c_i_* = *μ_c_* + *c_k_I_k_* + *σ_c_ϵ_c,i_*.

| Genotype | $r_i$ | $c_{kin}$ | $c_{non-kin}$ |
| --- | --- | --- | --- |
| A_05 | 2.13400444 | 1.154482646 | 1.215617111 |
| A_08 | 2.392899826 | 0.949480909 | 1.010615374 |
| A_11 | 2.391437999 | 1.533059983 | 1.594194448 |
| A_14 | 2.52212133 | 0.75051355 | 0.811648015 |
| F_07 | 1.833347256 | 1.090035607 | 1.151170072 |
| F_13 | 2.012928079 | 0.229535013 | 0.290669478 |
| F_15 | 1.902402391 | 1.369858669 | 1.430993134 |
| C_02 | 2.329240828 | 1.228480119 | 1.289614584 |
| M_12 | 2.524015276 | 1.617742492 | 1.678876957 |

Initial growth rates (*r_i_*) varied greatly among genotypes, but generally Algeria provenances had the fastest initial growth, followed by Morocco, Cyprus and France (Table 3). The growth of neighbor plants had a strong effect on the growth of focal genotypes (*c_i_*) for all the genotypes considered (*μ_c_* median value 1.06, credibility interval: 0.49; 1.913). The effect of *c_i_* was weaker in kin compared to non-kin mini-communities (median value for *c_k_* −0.06, credibility interval: −0.56;0.22) (Table 2). This indicates that the effect of neighbor plants’ growth on the growth of the focal plants was comparatively stronger in non-kin mini-communities. Initial growth *r_i_* (Table 3) had a strong effect on the growth of single genotypes (*μ_r_* median value 2.28, credibility interval: 2.03; 2.51).

To create a qualitative visual representation of individual genotypes’ growth as a function of initial growth rate *r_i_* and response to neighbors growth *c_i_*, we divided the genotypes into four different groups (marked by the median values of *r_i_* and *c_i_*) that could compare to the four different strategies predicted initially (Fig. 1a). While this representation is absolutely qualitative, it offers a rough categorization of genotypes with similar growth responses, and illustrates the absence of correlation between *c_i_* and *r_i_* (Fig. 1b).

### Growth and fruit production of focal plants

Overall, Focal plants grew bigger (during the first five weeks of the experiment) in non-kin mini-communities compared to kin mini-communities (Table 4). In average this difference in focal growth between community types was larger in high water treatments (13% difference in growth), compared to low water treatments (2% difference in growth) as shown by the marginally significant interaction between water treatment and community type (Table 4).

**Table 4:** results of linear models on leaves growth of focal plants of *Medicago truncatula*. Leaves growth was tested in response to community type (kin vs. non-kin), water treatment and regions of origin

| term | numDF | denDF | F- value | P-value |
| --- | --- | --- | --- | --- |
| CommunityType | 1 | 138 | 3.79 | 0.054 |
| WaterLevel | 1 | 138 | 18.32 | <0.001 |
| Origin | 2 | 138 | 0.78 | 0.460 |
| CommunityType:WaterLevel | 1 | 138 | 2.7 | 0.103 |
| CommunityType:Origin | 2 | 138 | 3.34 | 0.038 |
| WaterLevel:Origin | 2 | 138 | 0.62 | 0.542 |
| CommunityType:WaterLevel:Origin | 2 | 138 | 0.04 | 0.959 |

The growth response of focal plants to community types varied across origins (Table 4, Figure 2). Focal plants from Cyprus-Morocco grew in average 10% bigger in kin mini-communities compared to non-kin mini-communities. Focal plants from Algeria grew in average 7% larger in non-kin mini-communities compared to kin mini-communities. Focal plants from France, in average grew 16% bigger in non-kin compared to kin mini-communities. Post-hoc tests indicated significant differences between kin and non-kin mini-communities for France origins only (df= 137, t ratio = −2.918, p-value = 0.0041). This large variation in responses between kin and non-kin mini-communities was also found when visually inspecting genotypes combinations (Supporting information, Fig. S4). Models applied to radial growth rate as a function of origin, water treatment and community type produced analogous results (results not shown).

**Figure 2:**
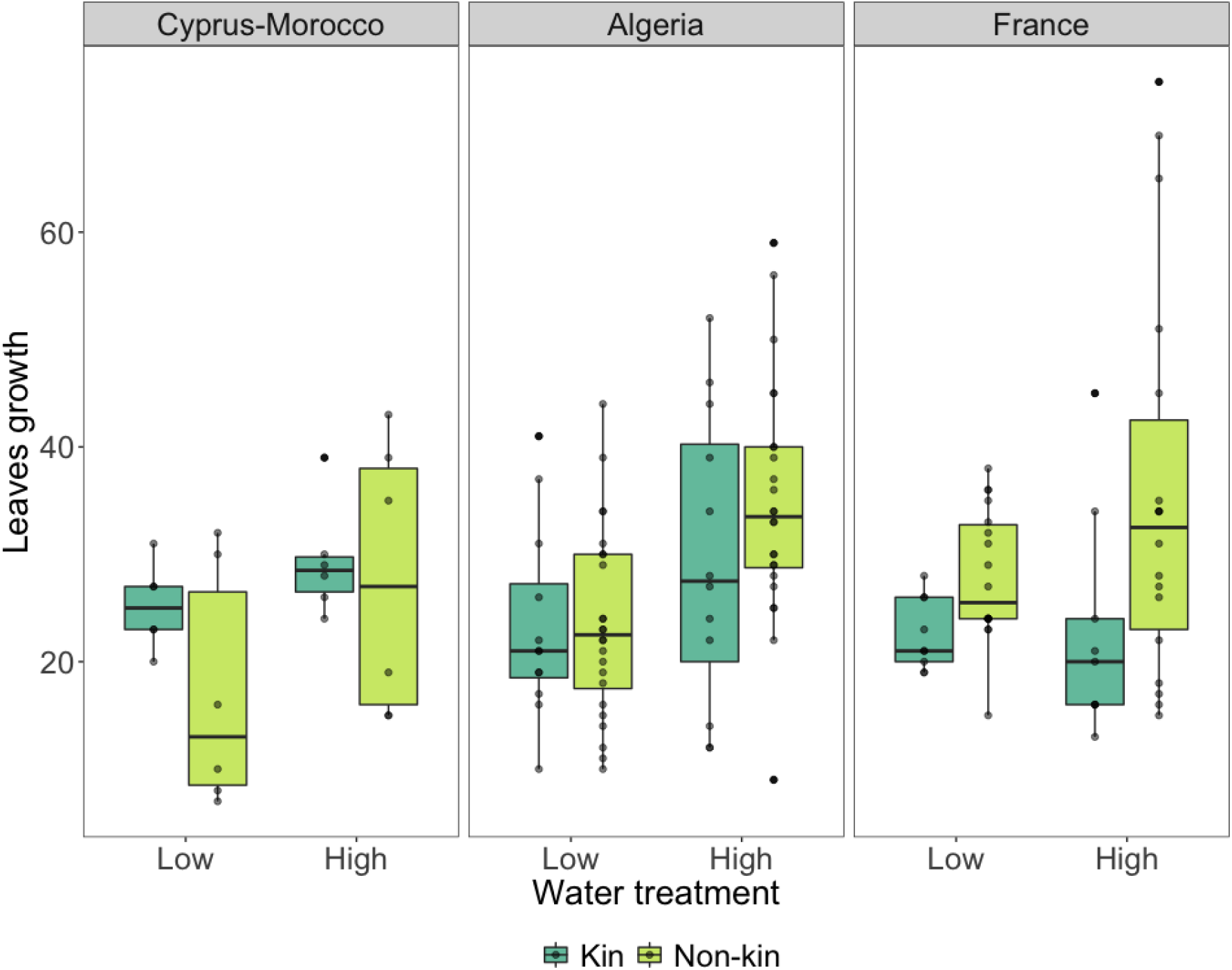
Boxplot representing leaves growth of focal plants as a response to community type and water treatment, across the different origins.

When looking at fruit production, we found a strong effect of water treatment, indicating that in average fruit production was three times larger in high water treatments (Table 5, Figure 3, Supporting information – Fig. S5). We did not find an effect of community type (Table 5, Fig. 3), but we found differences in fruit production among origins, with origins from Algeria and Cyprus-Morocco producing more fruits compared to France origins, which produced very few (Fig. 3, Table 5).

**Table 5:**
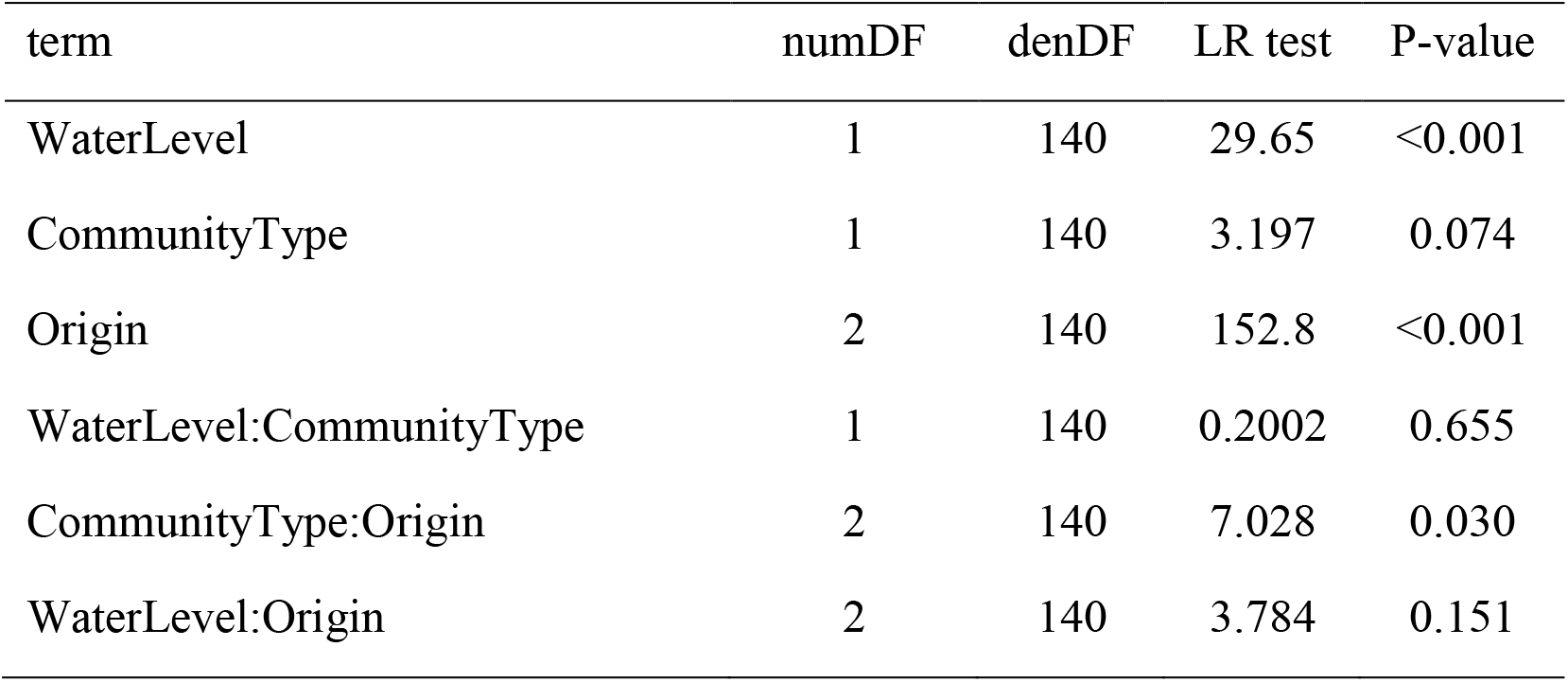
results of generalized models with negative binomial distribution on fruit production for *Medicago truncatula* in response to water treatment and focal genotype identity

**Figure 3:**
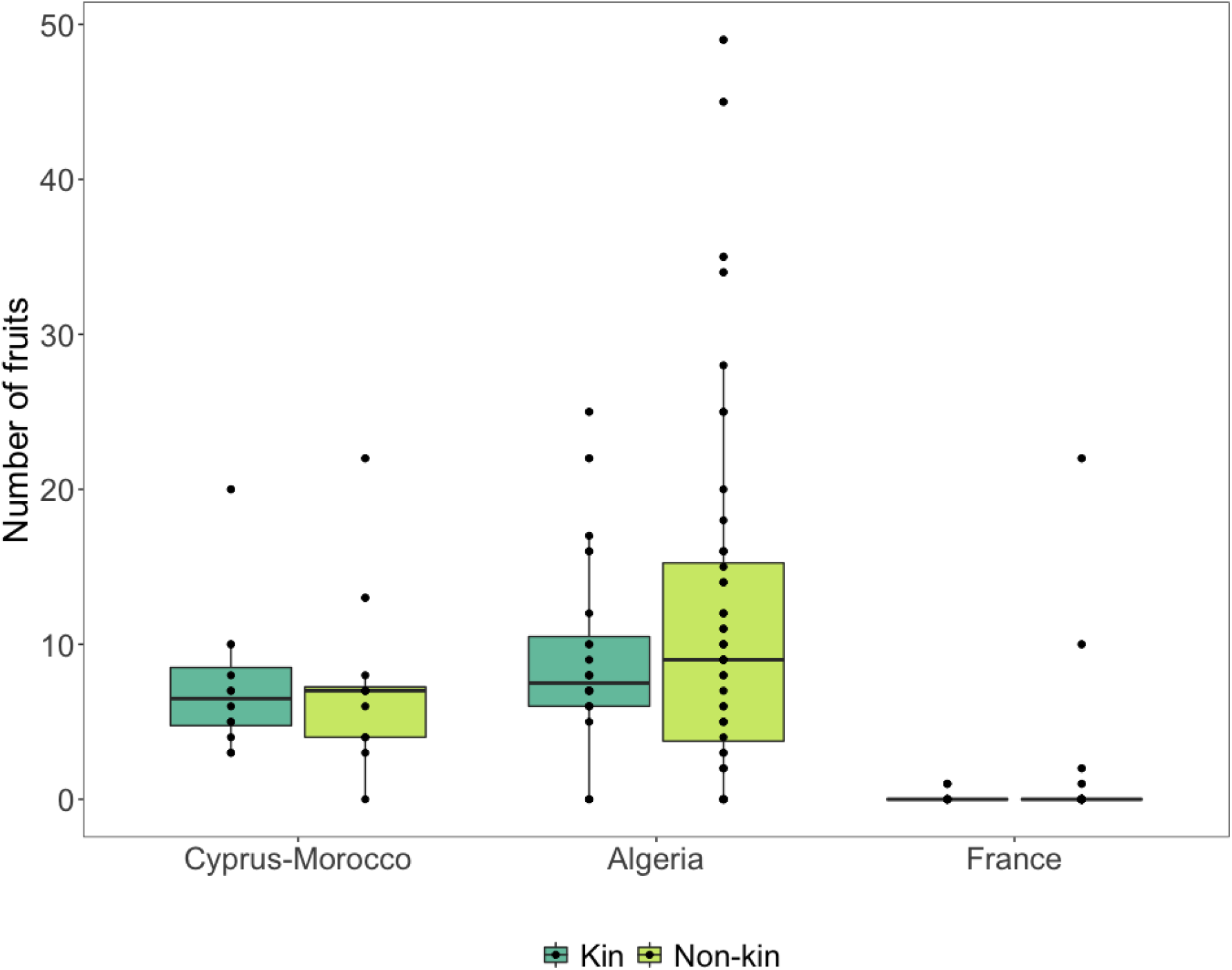
Boxplot representing fruit production of focal plants as a response to community type and origin.

When looking at mean fruit production for each genotype we found a correspondence between the genotypes that produced the highest number of fruits and those that, according to the Birch model, had the fastest initial growth rate (*r_i_*) and experienced less reduction in growth from neighbors (*c_i_*) (Supporting information - Fig. S5).

### Soil microbiome composition

16*S rRNA* amplicon sequencing yielded 5,461,606 reads distributed among the 66 samples, which were assigned to 1,585 microbial OTUs. Analysis of α-diversity (within sample diversity) revealed that soil in control pots (where no plants were growing) showed the highest α-diversity, compared with soil from either kin or non-kin mini-communities (Supporting information - Notes S2, Fig S6-S10). In addition, analysis of β-diversity using Principal Coordinate Analysis (PCoA) of weighted uniFrac distance showed that different genotypes had no significant influence on microbial composition (Fig. 4A). However, when comparing control pots with kin and non-kin mini-communities, we found a clear differentiation with different mini-communities explaining as much as 5.08% of the overall variance of 16*S rRNA* data.

**Figure 4:**
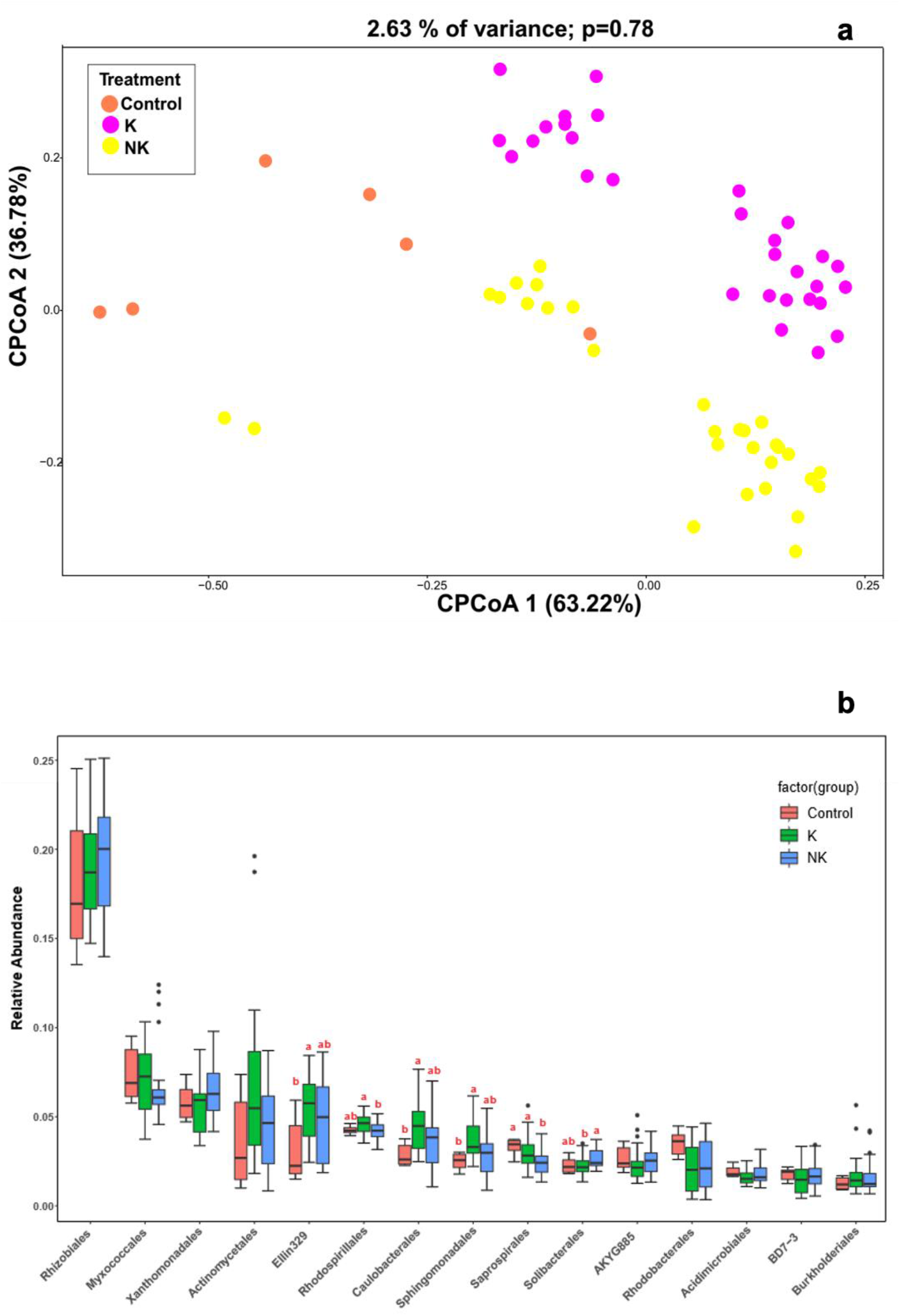
a) PCoA of microbial communities in control, kin and non-kin communities, where control represent treatments where no plants were grown in soil, b) Composition of soil microbial communities by taxa in control, kin and non-kin treatments, different letters represent significant differences across treatments based on Tukey HSD tests

Analyses of the 15 most abundant taxa among control, kin and non-kin mini-communities indicated that the abundance of Ellin329, Caulobacteriales and Sphingomonadales was significantly higher in kin mini-communities compared to control, whereas Saprospirales decreased significantly in non-kin mini-communities compared to control. Finally, the abundance of Rhodospirillales and Saprospirales was significantly higher in kin compared to non-kin communities and vice-versa Solibacterales were significantly more abundant in non-kin compared to kin communities (Fig. 5B). Differences among origins were negligible (Supporting information - Fig. S10).

## Discussion

We set out to investigate how the outcome of intraspecific interactions is mediated by genetic relatedness and water availability, and whether genotypic composition of mini-communities corresponds to different soil microbial communities. We used known concepts from species coexistence theory and from studies on intraspecific interactions to formulate hypotheses of how plants interact at the intraspecific level.

### Effects of relatedness

We predicted that in mini-communities of conspecifics, if genetic relatedness is a proxy for niche similarity, plant growth may be reduced by limited resources among kin, and influenced by genotype-specific traits among non-kin. We found that during early life stages focal plants grew larger in non-kin mini-communities compared to kin mini-communities (Fig. 2). This suggests that either plants growing with non-kin benefited from niche partitioning or that plants growing with kin may reduce growth toward their neighbors to avoid competition. Alternatively, if there is no reduced competition among kin, the lower biomass of focal plants in kin mini-communities may be explained by higher investment in root biomass (which was not measured in this experiment) leading to a lower aboveground biomass production. Notably, the growth response of focal genotypes to different mini-communities was different across origins. In our experiment we compared genotypes from France, where the maximum distance among genotype origins was approximately 30 km, from Algeria, where the spatial distance among origins ranged between 76 km and 490 km, and from Cyprus and Morocco, where genotypes origins were over 2400 km apart (Fig. S1). A significant difference in growth of focal plants among mini-communities (kin *vs*. non-kin) was found only in the case of France origins, suggesting that the effect of mini-community type was stronger for genotypes originating from similar sites, and that genetic relatedness may not the best predictor for the outcome of interactions among conspecifics. This result, is consistent with studies at the interspecific level, indicating that even among species, phylogenetic relatedness is not a reliable predictor for the outcome of interspecific interactions (Cahill Jr *et al*., 2008; Fritschie *et al*., 2014; Lyu *et al*., 2017).

The results of the Birch model, based on the estimates of final plant size at maturity indicated that, opposite to what we saw during early life stages, the effect of neighbors on the growth of focal genotypes (*c_i_*) was stronger in non-kin mini-communities, suggesting a stronger negative effect of competition among non-kin. This result may also be the consequence of larger differences in growth among non-kin mini-communities (as shown by the parameter *a_k_*) compared to kin mini-communities, due to the fact that identical genotypes share similar phenotypes and growth rates (Simonsen *et al*., 2014). However, even more interestingly, and somewhat consistently with the results relative to early life stages, the effects of mini-community type were weaker compared to those of genotype-specific initial growth rate (*μ_r_*) and response to neighbors’ growth (*μ_c_*). This indicates that genotype-specific traits, such as growth and response to competition may be comparatively more important than the genetic relatedness in determining focal plants growth response. Finally, when we plotted the values of initial growth rate (*r_i_*) and response to the growth of neighbors (*c_i_*), we found no relationship between the two parameters, potentially implying that growth rate and effect of competition from neighbors are unrelated (Fig. 1). The stronger effect of genotype-specific traits compared to community type detected in the Birch model is also consistent with the results on fruit production, which was driven mainly by water treatment and origin. Genotype-specific differences in fruit production (Fig. S5) were consistent (upon visual inspection) with the plot of Birch model estimates *r_i_* and *c_i_*. Overall, these results suggest that genetic relatedness among conspecific neighbors may have a stronger effect on plant growth (especially in early life stages) than on fitness, as shown by previous results (Milla *et al*., 2009, 2012). However, such effect seems to be limited to genotypes from close origins, and weaker compared to genotype-specific traits in affecting plant growth and fitness.

### Effects of environmental stress

We predicted that abiotic stress would mediate the outcome of plant-plant interactions across mini-community types. We used two different water treatments as a stressor because our target species *M. truncatula* is native to the Mediterranean region, where droughts are expected to become harsher according to climatic predictions (Rajczak & Schär, 2017). The results of the Birch model indicated, predictably, that the high water treatment greatly favored the total growth of plants at the pot level (*w_w_*, Table 1) and was also associated with lower degree of within-pot growth asymmetry (*a_w_*, Table 1). This was surprising because, according to the Stress Gradient Hypothesis (Bertness & Callaway, 1994), one would expect competition and possibly asymmetry in growth among individuals to increase under lower abiotic stress. It also has to be said that the effect of water treatment on asymmetric growth was smaller compared to the effect of community type (*a_k_*, Table 1), thus suggesting that growth asymmetry was more strongly influenced by differences in growth among genotypes than by water availability.

Water treatments mediated (though weakly) the effect of mini-community (kin *vs*. non-kin) type on focal plants’ leaves growth during the initial weeks. We found higher differences in growth of focal plants across communities in high water treatments compared to low water treatments. Interestingly, water treatments had a smaller effect on the growth of focal plants in kin mini-communities, compared to non-kin mini-communities (Fig.2). This suggests that, even in high availability of resources, kin may reduce growth to avoid competition among siblings, whereas in non-kin treatments focal growth may be largely influenced by growth rates of of the interacting genotypes,as predicted by our initial hypothesis. One reason for the relatively weak effect of water treatments in mediating the growth response across community types is that, despite most origins originated from similar climatic sites, the differences in drought extremes at each origin site may have had an impact on the effectiveness of the drought treatment (Supporting information - Fig. S4). Although, we observed signs of drought stress on the plants exposed to drought treatments, differences in drought tolerance among interacting genotypes may have added extra variation to our data.

### Soil microbial communities

We predicted that different genetic composition of our mini-communities may lead to different composition of soil microbial communities. At the interspecific level, soil microbial communities have been hypothesized to affect community coexistence (Bever *et al*., 1997) and several studies found indication of such patters between and within species (Laliberté *et al*., 2015; Keller & Lau, 2018; Siefert *et al*., 2018, 2019). We found the largest differences between the soil microbial of control pots, where no plants were grown, and the communities from kin and non-kin communities. However, we did not find large differences across kin and non-kin mini-communities in terms of soil community richness or diversity. Thus, these findings did not support our initial prediction that different genotypes may correspond to different composition of soil communities, and that this may reflect differences in plant growth and fitness. Notably, we had selected for analysis the soil samples corresponding to mini-communities where we observed the largest differences in plant growth between kin and non-kin. Our results are consistent with a previous study on *M. truncatula*, showing a low degree of differentiation in mycorrhizal colonization across its range (Dreher *et al*., 2017), and a long-term study of eleven species where increasing genotype diversity did not correspond to an increase of fungal community richness (Johnson *et al*., 2010). On the other hand, several studies have found a strong effect of plant genotype identity on soil community composition (Schweitzer *et al*., 2008; Micallef *et al*., 2009; Gehring *et al*., 2014) and plant-soil feedback (Semchenko *et al*., 2017).

One potential reason for the discrepancies among these studies and ours is the sampling method. We sampled soil from the center of the pot to have a soil sample that was representative of the whole soil community within a pot. However, this may have diluted the potential effect of each genotype on the soil microbiome. Had we also taken soil samples from the rhyzosphere and from the root surface, our results may have been substantially different. This indicates that single genotypes do not affect the composition of bulk soil, but do not exclude that single plants may have an impact of the composition of the rhizosphere microbiome.

### Final remarks

We found that intraspecific interactions in an annual legume are affected by the identity of neighbors (kin *vs*. non-kin), but this effect is stronger for genotypes from geographically close origins. Our results are consistent with both niche partitioning among non-kin and reduced competition among kin. The outcome of intraspecific interactions was also mediated by water availability. While the response of focal plants to kin neighbors did not change water treatments, suggesting the plants may reduced their growth to avoid competition among kin, higher water treatments corresponded in average to higher growth among non-kin, suggesting that in this case plant growth may be mediated by individual genotypes growth rates, as also predicted by our model. we found that in non-kin treatments This is consistent with previous model on species coexistence that stress how species coexistence may be mediated rather than by niche differentiation, by variation in competitive ability among genotypes within species (Begon & Wall, 1987) and with empirical studies (Cahill *et al*., 2005, and @masclaux_competitive_2010; Fridley & Grime, 2010).

Interestingly, the effects of neighborhood were limited to plant growth, but did not influence fruit production. Lastly, drought stress exacerbated differences in performance among communities, indicating that interactions among kin may be governed by competition avoidance, and interactions among non-kin may be largely affected by genotype-specific traits and resource availability. However, these effects should be tested in a broader context, accounting for different drought tolerances among origins. Our study, indicates a strong intraspecific variation in M. truncatula and a potential for genotypic variation in traits to play and important role in intra- and inter-specific interactions.

## Acknowledgments

We thank L. Lauridsen and J. Rytter for assistance in the greenhouse. Pierre Liancourt contributed insightful comments and suggestions for the conceptual development of the manuscript. This work was funded by a grant from the Danish Council of Independent Research (grant number 6108-00200B) to BKE.

## Author contributions

ST, BKE, CFD and SR planned the experiment, JMP, LG JR provided plant material, ST carried out the experiment and sampling, ST and KT sampled soil, ST, SK and SZ conducted DNA extraction on soil samples, KT and SZ conducted bioinformatics analyses and multivariate statistics on soil samples, ST wrote the first draft of the ms and all authors contributed to successive versions.

## Data availability

Data will be made available in a data repository upon acceptance of the manuscript

## Supporting information

**Figure S1** Map of the origin sites of the *Medicago truncatula* genotypes used for this experiment. Distance between sites ranges between 0 and 30 km in France, and between 120km and 450 km in Algeria.

**Figure S2** Plot showing the relationship between spatial distance among sites of origin of the genotypes (measure in km) and their genetic relatedness (calculated using the Rousset index).

**Figure S3** Estimate of volume in leaves equivalent based on regression of plant volume (calculated as *πr^2^h*) and early measurements of leaves number.

**Figure S4** Boxplot showing leaves growth (measured as increment in number of leaves) in response to community type and water treatment across single genotypes.

**Figure S5** Boxplot showing fruit production (measured as number of fruits) in response to community type and water treatment across single genotypes.

**Figure S6** Boxplot representing the Shannon Index calculated for soil microbial communities.

**Figure S7** Boxplot showing relative abundance of microbial taxa found in soil communities of a) Cyprus (genotype C002) and b) Morocco (genotype M012).

**Figure S8** Boxplot showing relative abundance of microbial taxa found in soil communities of Algeria origins for a) genotype A008 and b) genotype A014.

**Figure S9** Boxplot showing relative abundance of microbial taxa found in soil communities of France origins (genotype F015).

**Figure S10** Barplot showing relative abundance of microbial species for each soil sample analyzed in this study.

**Table S1** List of *Medicago truncatula* genotypes used for the experiment with corresponding accession labels, Hapmad ID and origins site coordinates used for the experiment.

**Table S2** Altitude (measured in m asl) and climatic variables for the origin sites of each genotype of *Medicago truncatula* used in this experiment.

**Table S3**: list and explanation of the parameters used in the Birch model.

**Notes S1** detailed description of the growth model

**Notes 2** Detailed results for soil microbial communities

