## Supplementary_Information_files for "Identity and provenance of neighbors, genotype-specific traits and abiotic stress affect intraspecific interactions in the annual legume *Medicago truncatula*"

**The following supporting information is available for this manuscript**

**Table S1: list of *Medicago truncatula* genotypes used for the experiment with corresponding accession labels, Hapmad ID and origins site coordinates used for the experiment**

| Genotype | Accession label | Hapmap_ID | Latitude | Longitude |
|---|---|---|---|---|
| A_05 | DZA315-16 | HM005 | 34.716N | 0.158E |
| A_08 | DZA012-J | HM008 | 36.549N | 3.183E |
| A_11 | DZA-327-7 | HM011 | 35.252N | 0.703W |
| A_14 | DZA233-4 | HM014 | 35.847N | 4.951E |
| F_07 | Salse071B | HM007 | 42.820N | 2.945E |
| F_13 | Salse042B | HM013 | 42.820N | 2.945E |
| F_15 | F11013-3 | HM015 | 43.090N | 2.86299988746643E |
| C_02 | SA028064TR.1 | HM002 | 34.783N | 33.167E |
| M_12 | SA026063TR.1 | HM012 | 32.167N | 8.833W |

**Figure S1: map of the origin sites of the *Medicago truncatula* genotypes used for this experiment. Distance between sites ranges between 0 and 30 km in France, and between 120km and 450 km in Algeria.**

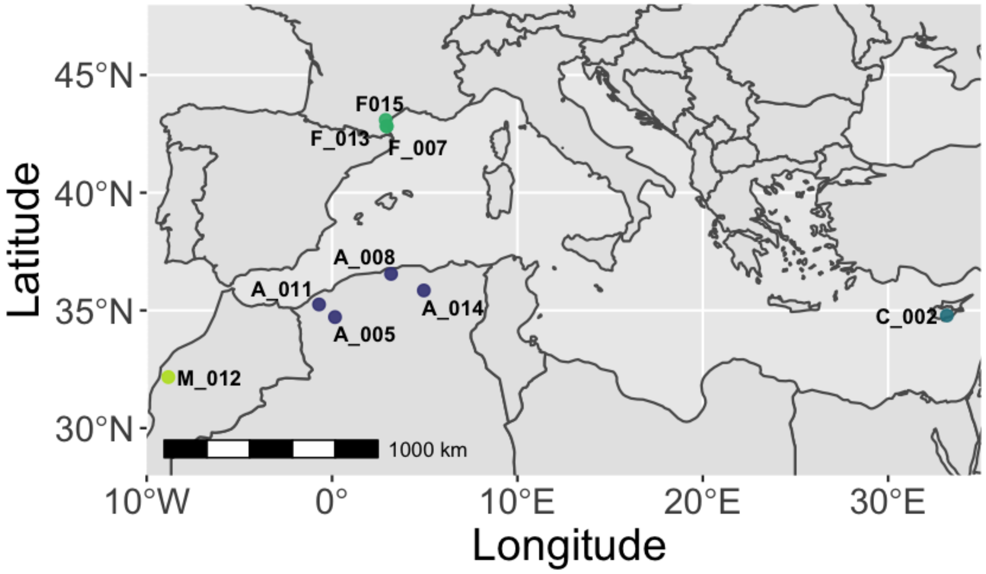

**Table S2: Altitude (measured in m asl) and climatic variables for the origin sites of each genotype of *Medicago truncatula* used in this experiment (data from Burgarella *et al.* (2016)). mean T wettest and mean T driest define the mean temperature (measured in Celsius) of the wettest quarter and the driest quarter respectively; annual precipitation, precipitation driest (i.e. precipitation of the driest quarter) precipitation wettest (i.e. precipiation of the wettest quarter) are measured in mm.**

| Genotype | Altitude (m asl) | mean T wettest | mean T driest | annual precipitation | precipitation driest | precipitation wettest |
|---|---|---|---|---|---|---|
| A_05 | 1098 | 7.51 | 24.1 | 345 | 40 | 112 |
| A_08 | 266 | 11.63 | 23.4 | 699 | 26 | 326 |
| A_11 | 490 | 9.95 | 23.1 | 457 | 20 | 199 |
| A_14 | 1383 | 4.45 | 21.65 | 471 | 48 | 156 |
| F_07 | 6 | 12.28 | 22.8 | 587 | 88 | 200 |
| F_13 | 6 | 12.28 | 22.8 | 587 | 88 | 200 |
| F_15 | 111 | 11.13 | 21.83 | 653 | 105 | 210 |
| C_02 | 258 | 10.58 | 24.81 | 504 | 8 | 309 |
| M_12 | 358 | 13.05 | 24.25 | 336 | 8 | 148 |

**Figure S2: plot showing the relationship between spatial distance among sites of origin of the genotypes (measure in km) and their genetic relatedness (calculated using the Rousset index). Only genotype combinations used for the experiment are represented.**

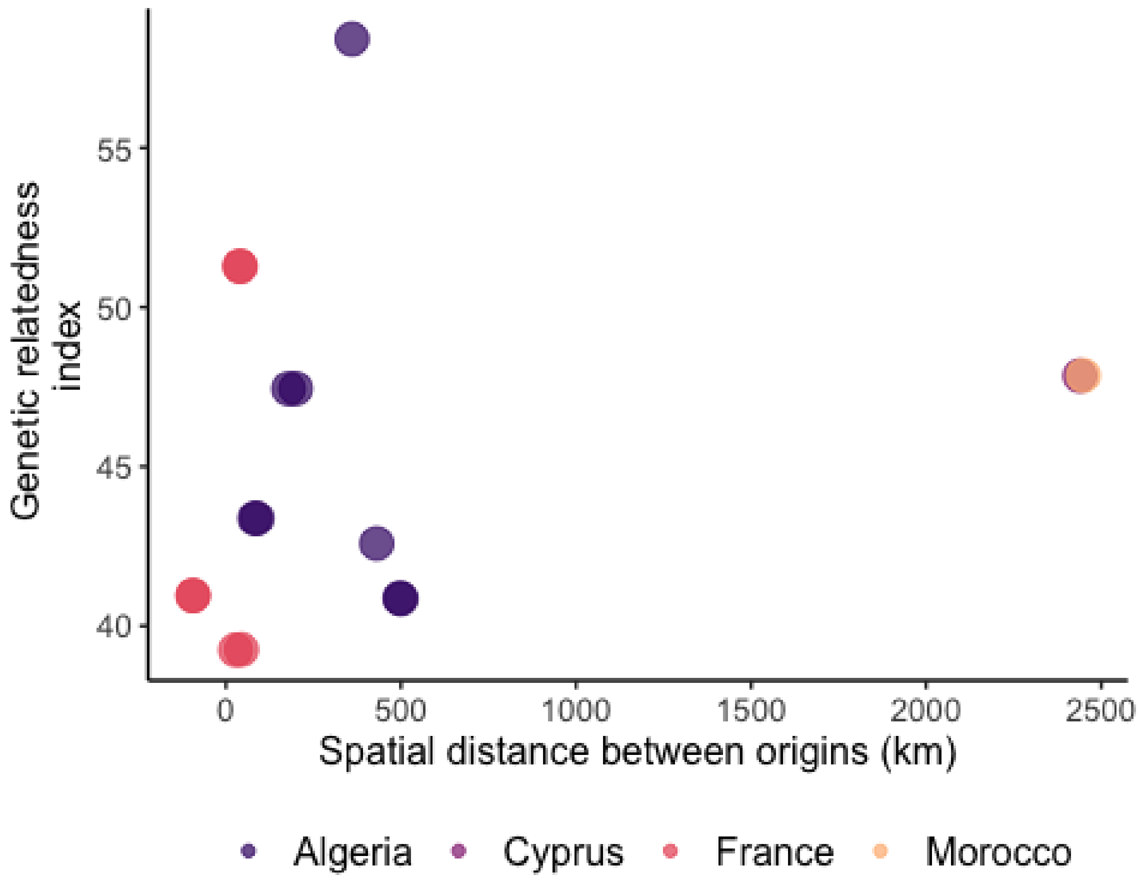

**Figure S3: estimate of volume in leaves equivalent based on regression of plant volume (calculated as $\pi r^2 h$) and early measurements of leaves number.**

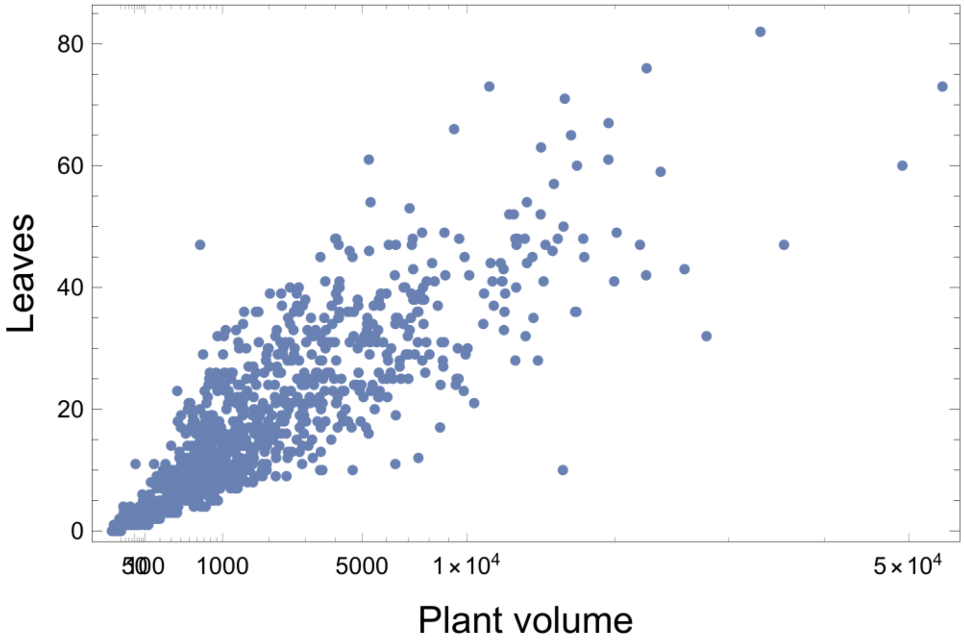

**Notes S1: detailed description of the growth model**

The generalized growth model, initially described in Damgaard & Weiner (2008) , was based on a growth model proposed by Birch (1999) :

$$\frac{dv(t)}{dt} = rv(t)\frac{w - v(t)}{w - v(t + cv(t))} (Eqn. 1)$$

where

$$v \geq 0, r > 0, c \geq 0$$

As detailed in Damgaard & Weiner (2008), *v(t)* is the plant size at time *t*, *r* is the initial relative growth rate, *w* is the final plant size when growth stops. The point at which *w* is reached is affected by *r* and by *c*

**The inflection point: c**

*c* is a positive parameter that determines the inflection point of the growth curve after reaching which, the asymptote is reached. This parameter describes the effect of the growth of neighbors on the growth of the focal. When this effect is stronger, the focal plant will reach the inflection point and stop growing earlier. Larger values of *c* indicate that the inflection point is reached earlier and that competition from neighbors begin to become important and reduce the growth of focal plant early on in its development.

In the Birch model, the growth of a plant or of a genotype is represented by an exponential curve when c = 0 and by logistic curve when c = 1. The maximum growth rate is $\frac{rw}{(\sqrt{c}+1)^2}$ when the plants have the size $\frac{w}{(\sqrt{c}+1)}$ that is, when $c < 1$, the maximum growth occurs at $v > w/2$; when $c > 1$ the maximum growth occurs at $v < w/2$.

**The size asymmetry coefficient: a**

The model also allows to take into account variations in individual growth rate and size that occur as a consequence of genotype identity, time of germination, microenvironment, etcetera. These differences in growth can become important when plant growth is limited by a resource that may be monopolized. In this case, asymmetric competition may occur leading to faster growth of larger plants at the expense of smaller ones resulting in size-asymmetric growth (Damgaard & Weiner (2008)).

The size asymmetry among individuals is described by a coefficient $a$. Negative values of $a$ indicate that the growth rate of a plant is less than proportional to its size, whereas positive values indicate that a plant's growth rate is more than proportional to its size.

Size-asymmetric growth can be integrated into the Birch growth model by assuming that individual plant growth is proportional to a power function of size (Schwinning & Fox, 1995; Damgaard, 1999; Wyszomirski *et al.*, 1999; Damgaard *et al.*, 2002).

(Eqn. 2)

$$f(v(t), a) = (v(t) + 1)^a - 1$$

with a>0

The relationship between plant size and growth rate is expressed with a size-asymmetry coefficient, *a*, which measures the degree of curvature of the growth-size relationship within the population over the entire growth curve. Here, negative values of *a* indicate that growth rate is less than proportional to the size of the plant, whereas positive values of *a* indicate that growth rate is more than proportional to plant size within a population (Schwinning & Weiner, 1998; Weiner & Damgaard, 2006).

The individual-based Birch growth model considers the effect of plant size variation on the growth of individual plants by generalizing (Eqn. 1) with respect to size-asymmetric growth (Eqn. 2). Assume a population of *n* competitively interacting plants of variable size, then the growth of plant *i* at time *t* may be expressed by *n* coupled differential equations (here described by a single equation for plant *i*),

(Eqn. 3)

$$\frac{dv\ i\ (t)}{dt} = \frac{r\ i\ (v\ i\ (t) + 1)^a - 1}{a} \frac{nw - \sum_{k=1}^{n} v_k\ (t)}{nw - (1 - c_i \sum_{k=1}^{n} v_k\ (t))}$$

with $v_i \geq 0, a \neq 0, r_i > 0, w > 0, c_i \geq 0$ where $v_i(t)$ is the size of plant *i* at time *t*.

with $v_i \geq 0, a \neq 0, r_i > 0, w > 0, c_i \geq 0$ where $v_i(t)$ is the size of plant *i* at time *t*.

**Genetic variation among growth curves: $r_i$ and $c_i$**

To include variation among growth curves, we allowed for variation among genotypes in the two parameters $r_i$ $c_i$, which determine respectively the initial relative growth rate, and the shape of the growth curve. The other two parameters, *a*, the size-asymmetry coefficient, and *w* the average plant size at the end of the growing season, were set to a fixed value for the entire population (in our case, a population is defined by the group of individuals within the same pot). Generally, larger plants grow more, and growth is reduced by large total population biomass (Damgaard & Weiner (2008)). Similarly to te Birch growth model, the initial relative growth rate of plants of genotype *i* in growth model (Eqn. 3) is equal to $r_i$ at the limit when all *n* plants are small, i.e.

$$\lim_{\forall v_k \to 0} \frac{1}{v_i(t)} \frac{dv_i(t)}{dt} = r_i$$

At the limit when a = 0 , model (3) is not defined but the growth curve is continuous in a = 0 with

$$\lim_{a \to 0} \frac{r_i (v_i(t) + 1)^a - 1}{a} = r_i log(v_i(t) + 1)$$

the size where the plant *i* experiences the maximum growth rate is the solution to $v_i''(t) = 0$ which may be solved numerically in the simplifying case when w = 1, $v_i = p$, and $\sum v_k \approx np$ where p is the relative size to w.

the size where the plant *i* experiences the maximum growth rate is the solution to $v_i''(t) = 0$ which may be solved numerically in the simplifying case when w = 1, $v_i = p$, and $\sum v_k \approx np$ where p is the relative size to w.

*Parameter estimation:*

To minimize auto-correlation among the residual variation, the growth model (Eq. 3) was fitted to the observed growth increments in plant size (Seber & Wild, 1989). The expected sizes of the plants at time *t* were calculated using the observed plant sizes at time *t-1* and a specific parameterization of the growth model and compared to the observed plant sizes at time *t*,

( Eq. 4)

$$V_i(t_i) = \int_{t_{j-1}}^{t_j} \frac{dV_i(t; V_i(t_{j-1}), a, w, r_i, c_i)}{dt} dt + \varepsilon_{ij}$$

with $i = 1,2,\dots, n$; $j = 2,3,\dots,m$

where *m* is the number of observed changes in plant size, $V_i(t_j)$ is the observed plant size at time $t_j$, $\frac{dV_i(t;V_i(t_{j-1}),a,w,r_i,c_i)}{dt}$ is the expected plant growth of plant *i* at time *t* calculated from (Eq. 3) using the NDSolve routine of Mathematica (Wolfram, 2003), and where $V(t_{j-1})$ is a vector of the observed plant sizes at time $t_{j-1}$.

The residual error was assumed to increase proportionally with expected plant size and the time period of growth, and modelled by a Student's t distribution, i.e., $\varepsilon_{ij}\ T(0, v, \sigma V_i(t_j)(t_j - t_{j-1}))$.

The joint Bayesian posterior distribution of the parameters were simulated by an MCMC approach using the Metropolis Hastings algorithm, assuming uniform improper prior distributions in the domains of the parameters. The MCMC iterations converged relatively fast and after a burn-in period of 10,000 iterations the next 40,000 iterations were to calculate the marginal posterior distribution and the corresponding 95% credibility interval of each parameter.

**Table S3: list and explanation of the parameters used in the Birch model**

| Parameter | Interpretation | dependent upon |
|---|---|---|
| *w* | final plant size at the pot level, | water treatment |
| *a* | within-pot size asymmetric growth | water treatment community type (kin vs. non-kin) |
| $r_i$ | initial growth of genotype *i* in the absence of competition | plant genotype |
| *c* | inflection point of the growth curve of genotype *i* where plant growth stabilizes | plant genotype identity, community type |

**Figure S4: Boxplot showing leaves growth (measured as increment in number of leaves) in response to community type and water treatment across single genotypes from Algeria (A_005, A_008, A_011, A_014), France (F_007, F_013, F_015), Cyprus (C_002) and Morocco (M_012). Semi-transparent dots represent focal individuals, full dots represent outliers.**

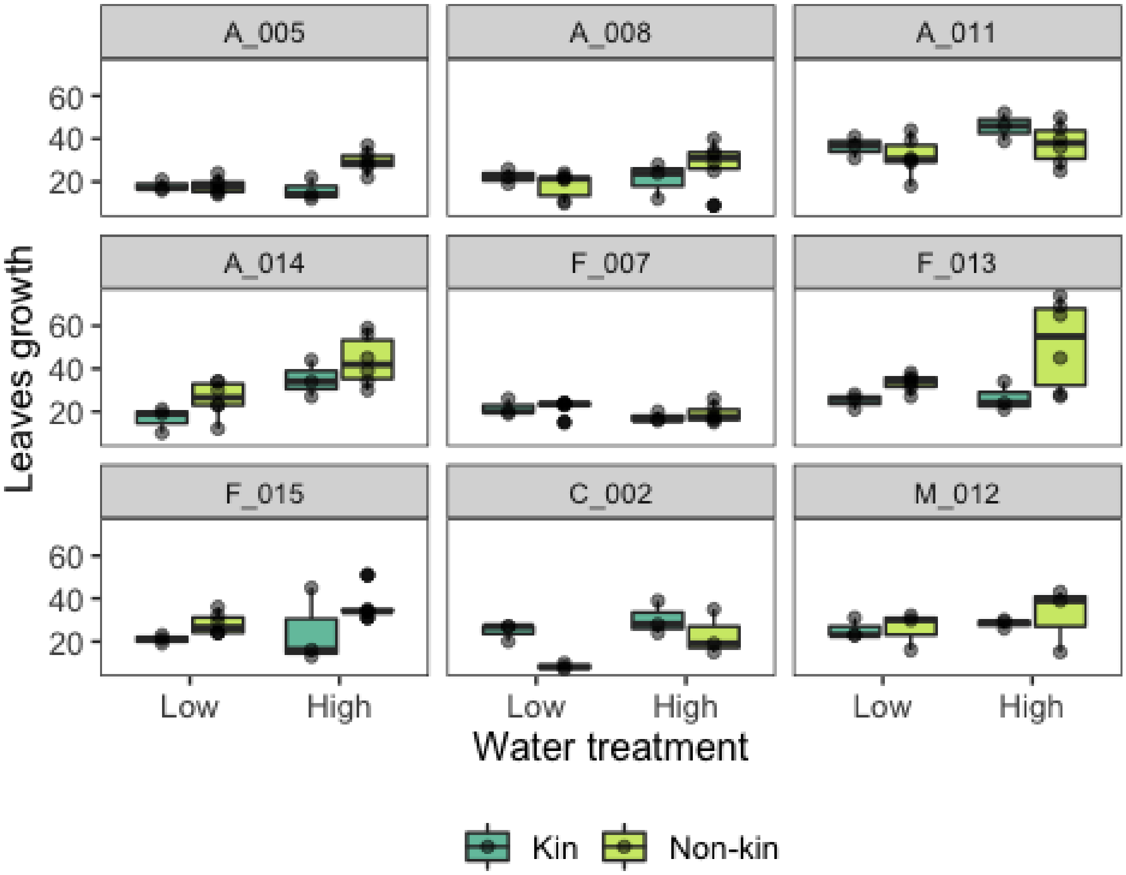

**Figure S5: Boxplot showing fruit production (measured as number of fruits) in response to community type and water treatment across single genotypes from Algeria (A_005, A_008, A_011, A_014), France (F_007, F_013, F_015), Cyprus (C_002) and Morocco (M_012). Semi-transparent dots represent focal individuals, full dots represent outliers.**

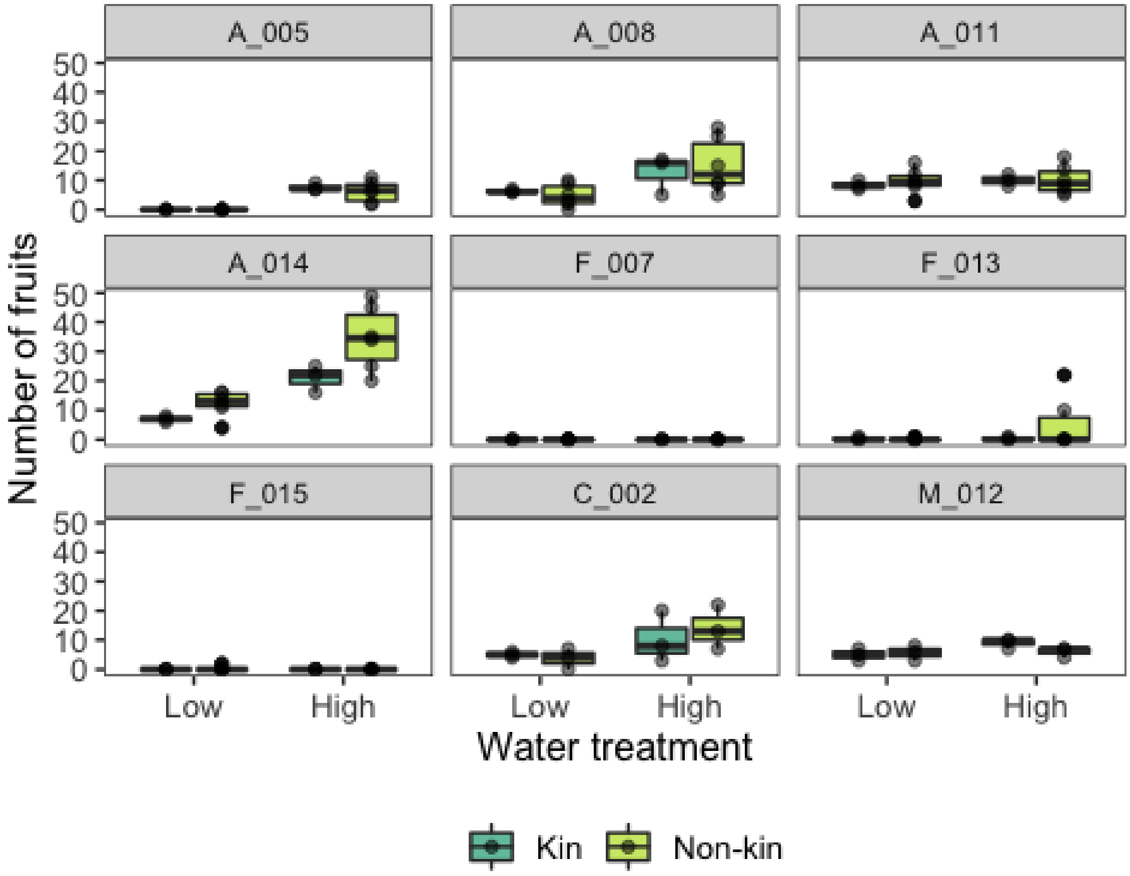

**Notes S2: Detailed results of soil microbial communities**

When comparing soil microbiome from kin and non-kin mini-communities (focal plants A08, A14, F15 and M12), we found no significant difference in α-diversity, with the only exception of focal genotype C2, which had significantly higher α-diversity in non-kin mini-communities (Fig. S6).

Analysis of taxonomic composition profile revealed high similarity of microbial community composition among soil samples from control pots and from mini-communities (kin and non-kin). This suggests that different plant genotypes have a minimal effect on the microbial composition at the whole community level.

When comparing the taxa in soil samples from control pots and from mini-communities, we found that in soil samples from mini-communities the abundance of Ellin329 and Caulobacteriales was significantly higher and the abundance of Saprospirales and Rhodobacterales was significantly lower compared to control pots (results not shown). When looking at each of the mini-communities separately, we found that in soil from Cyprus-Morocco mini-communities (focal genotype C2, Fig. S7-a) the relative abundance of Actinomycetales, Ellin329, Caulobacteriales and Sphingomonadales was higher in kin compared to non-kin mini-communities, while Rhodobacterales had a lower abundance in kin mini-communities. Additionally, in Cyprus-Morocco mini-communities (focal genotype M12, Fig. S7-b) the abundance of Actinomycetales, Sphingomonadales and Saprospirales was higher in kin compared to non-kin mini-communities. Conversely, Myxococcales more abundant in non-kin mini-communities (Fig. S8)

In the analyzed soil samples of mini-communities from Algeria origins (focal genotypes A_008, Fig. S8-a) we found higher abundance of Myxococcales and Rhodospirillales in kin compared to non-kin mini-communities. However, Solibacterales and Acidimicrobiales were less abundant in kin mini-communities. Xanthomonadales and Acidimicrobiales (focal genotype A14, Fig. S8-b) were more abundant in non-kin compared to kin communities.

Finally, soil samples of mini-communities from France origins (focal genotype F_015, Figure S9) did not show any significant difference in microbial taxa abundances between mini-communities

Comparison of relative abundance of the 15 most abundance microbial taxa, with low abundance taxa grouped on the bottom, does not show significant differences in the relative abundance across genotype combinations (Fig S10). Overall, our results do not indicate a large influence of plant genotype identity on soil microbiome. However, our analyses were limited to the soil between single plants, and soil samples from the rhizosphere may show a larger differentiation in the composition of soil biota.

**Figure S6: boxplot representing the Shannon Index calculated for soil microbial communities.**

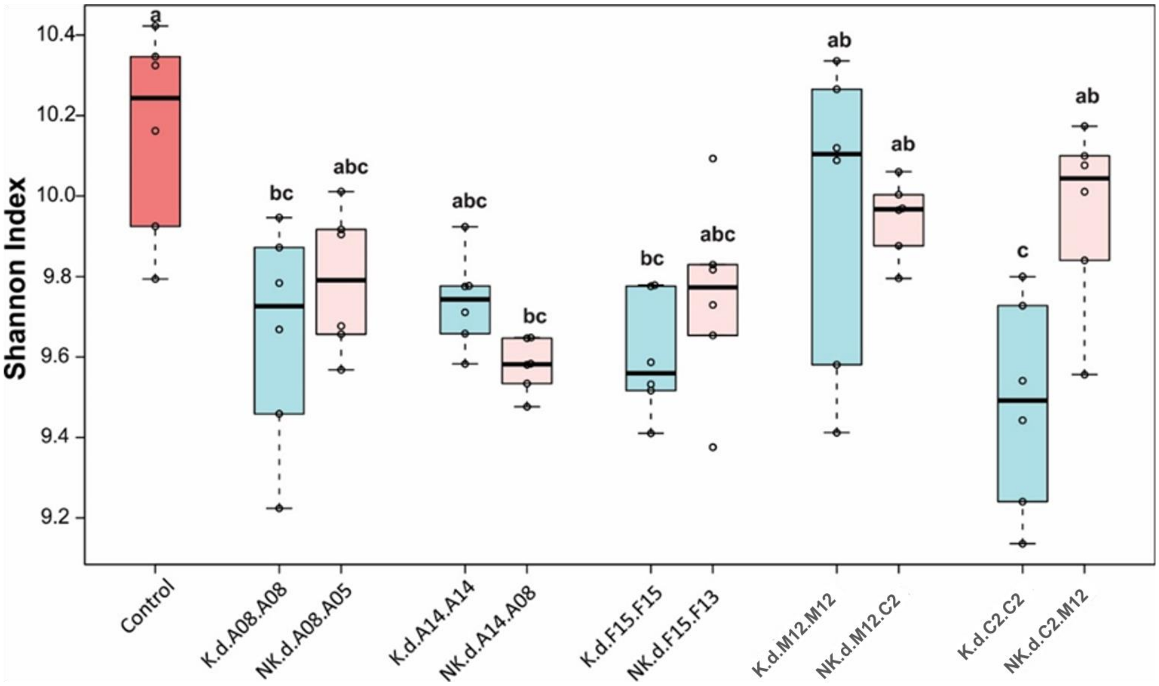

**Figure S7: boxplot showing relative abundance of microbial taxa found in soil communities of a) Cyprus (genotype C002) and b) Morocco (genotype M012).**

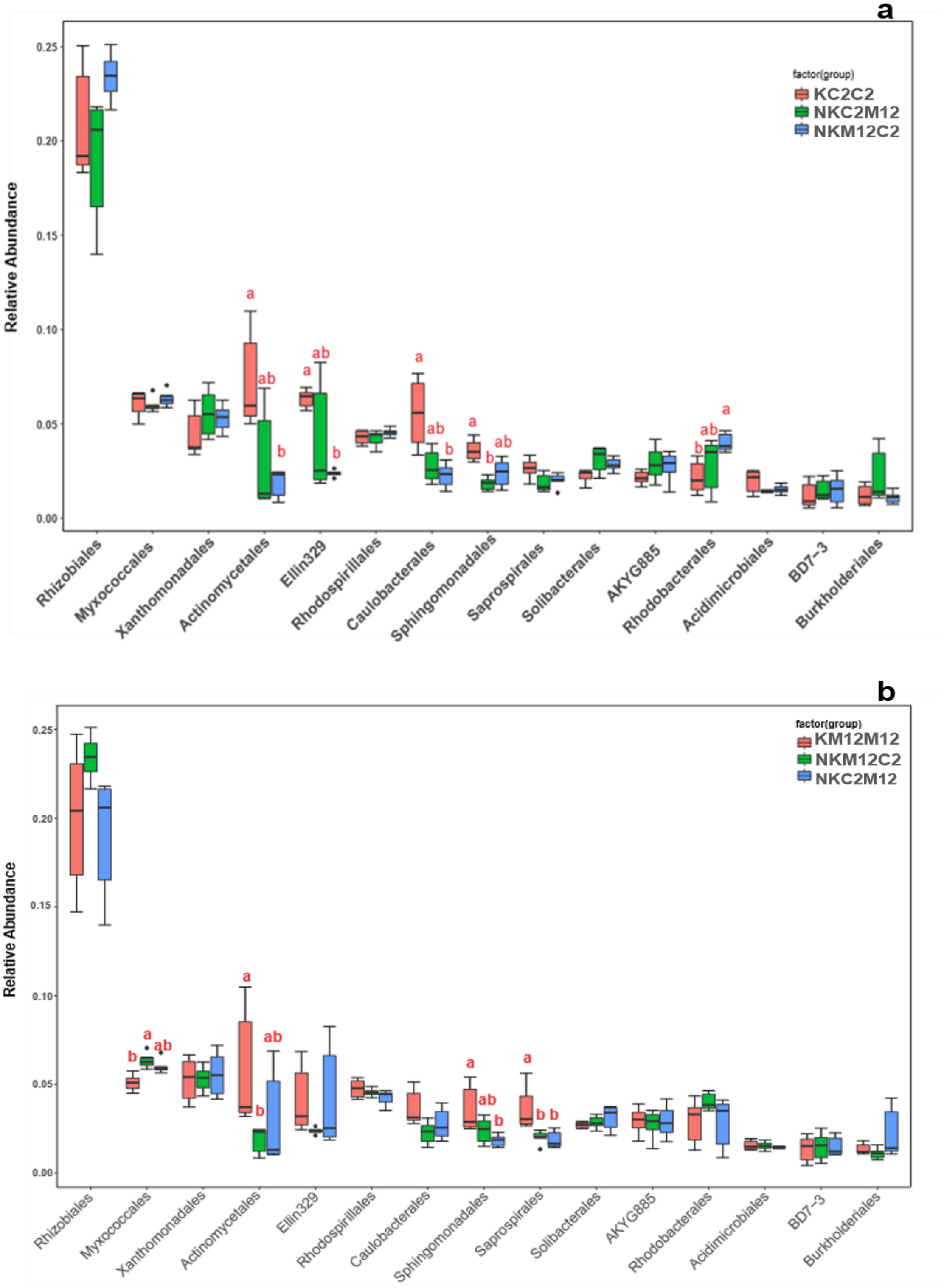

**Figure S8: boxplot showing relative abundance of microbial taxa found in soil communities of Algeria origins for a) genotype A008 and b) genotype A014.**

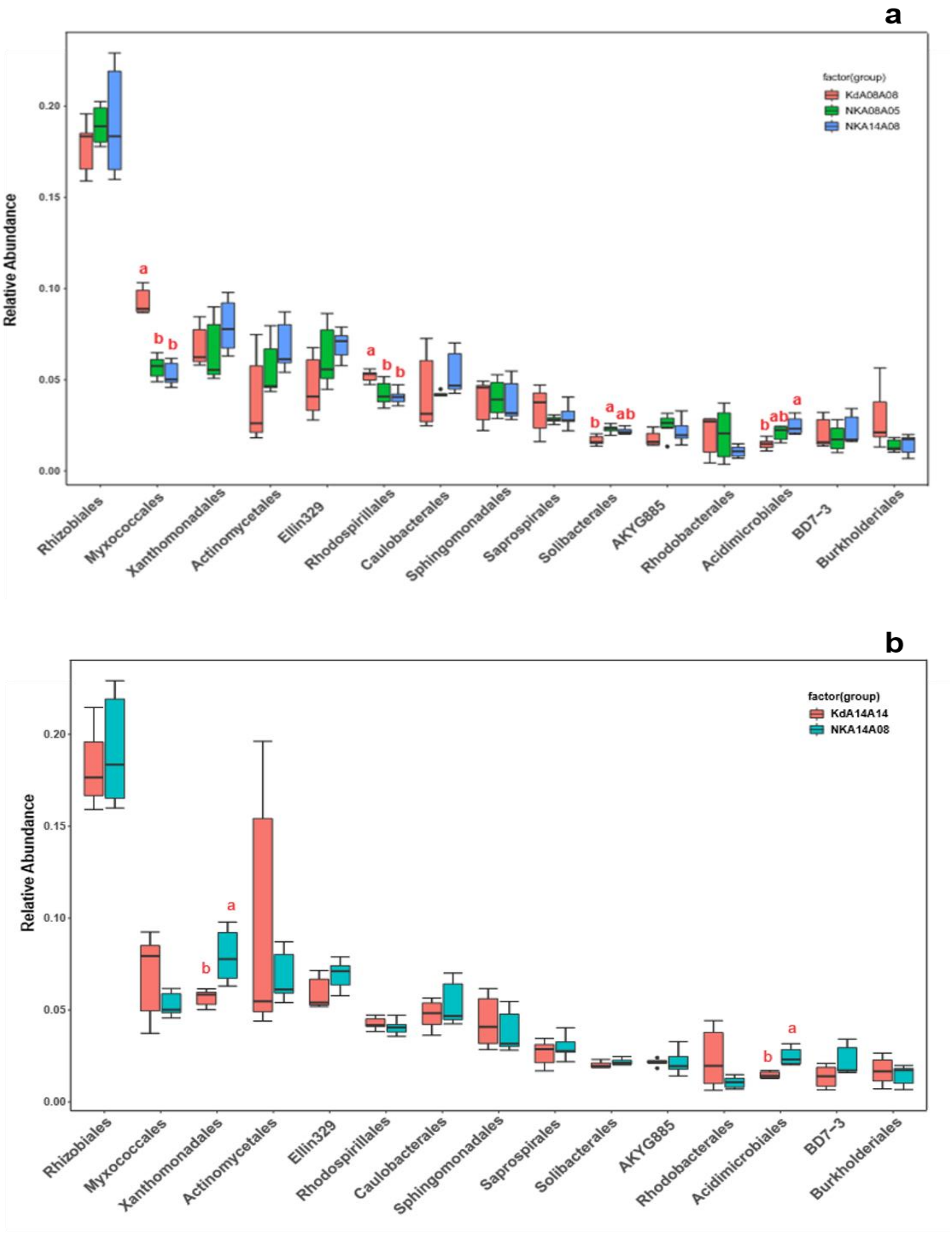

**Figure S9: boxplot showing relative abundance of microbial taxa found in soil communities of France origins (genotype F015).**

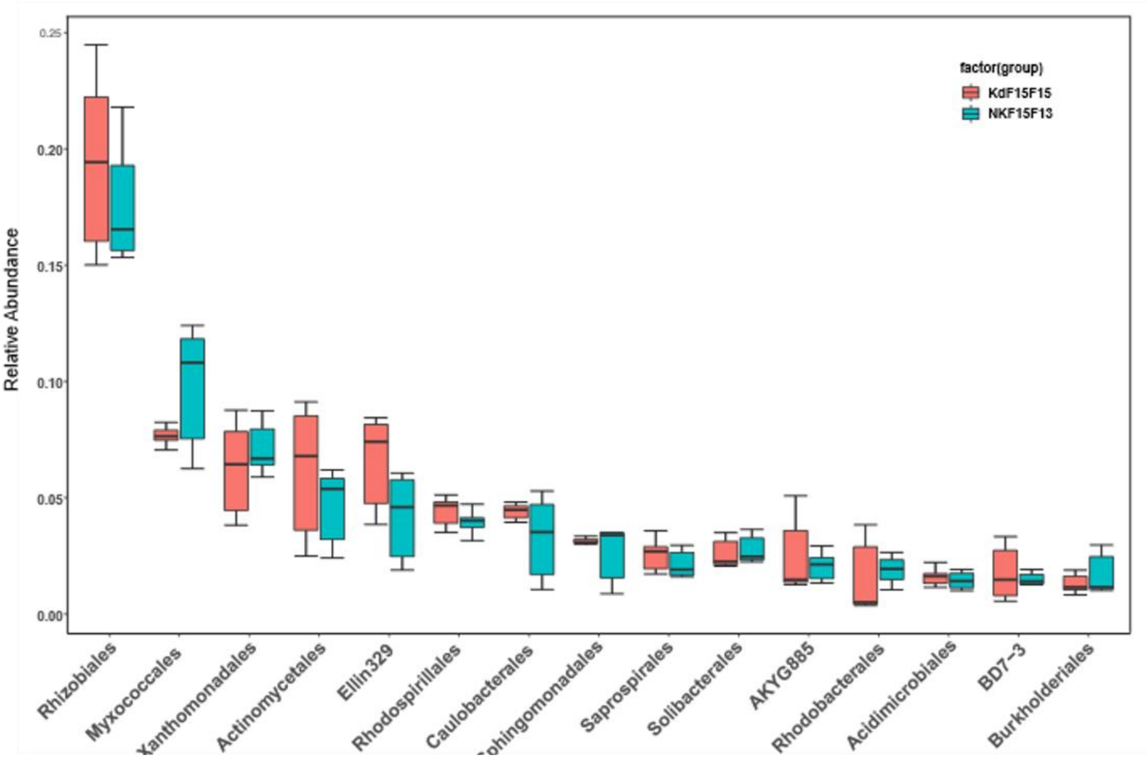

**Figure S10: Barplot showing relative abundance of microbial species for each soil sample analyzed in this study.**

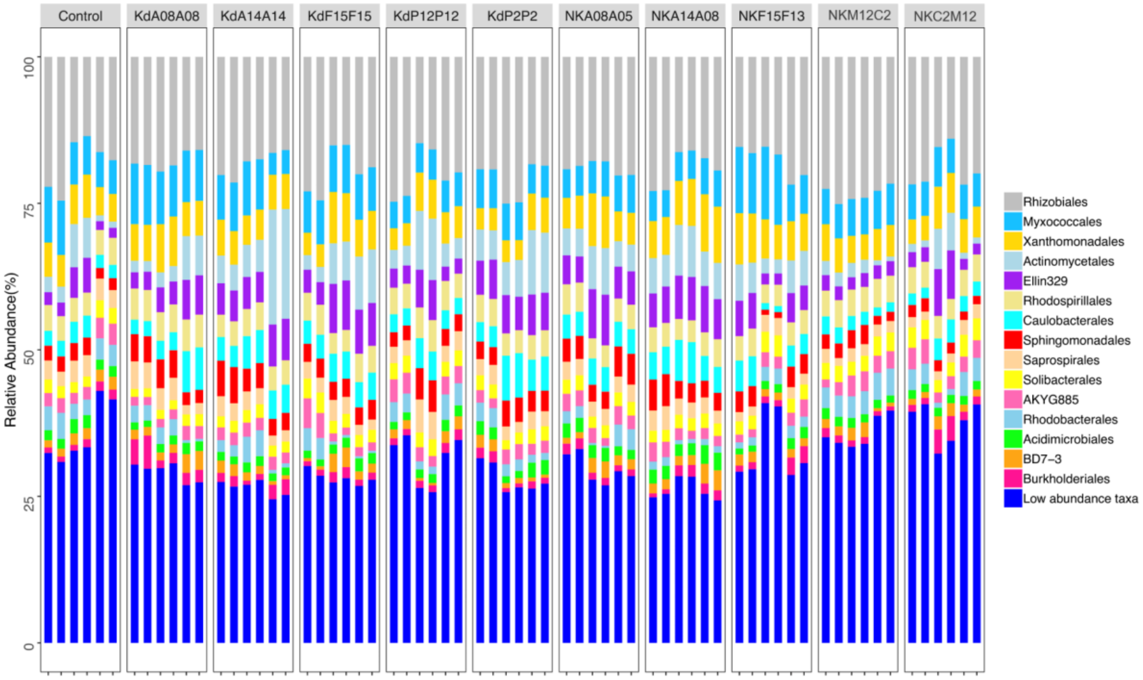
